# Alternative Splicing of MR1 regulates antigen presentation to MAIT cells

**DOI:** 10.1101/695296

**Authors:** Gitanjali A. Narayanan, Abhinav Nellore, Jessica G. Tran, Aneta H. Worley, Erin W. Meermeier, Elham Karamooz, Megan Huber, Regina Kurapova, Fikadu Tafesse, Melanie J. Harriff, David M. Lewinsohn

**Affiliations:** Department of Biomedical Engineering, Oregon Health and Science University, Portland, OR, USA; Department of Computational Biology, Oregon Health and Science University, Portland, OR, USA; VA Portland Health System, Portland, OR, USA; Department of Pulmonary and Critical Care Medicine, Oregon Health and Science University, Portland, OR, USA; Department of Molecular Microbiology and Immunology, Oregon Health and Science University, Portland, OR, USA

**Keywords:** Antigen presentation, T cell activation, Alternative splicing

## Abstract

Mucosal Associated Invariant T (MAIT) cells can sense intracellular infection by a broad array of pathogens. These cells are activated upon encountering microbial antigen(s) displayed by MR1 on the surface of an infected cell. Human MR1 undergoes alternative splicing. The full length isoform, MR1A, can activate MAIT cells, while the function of the isoforms, MR1B and MR1C, are not well characterized.

In this report, we sought to characterize these splice variants. Using a transcriptomic analysis in conjunction with qPCR, we find that that MR1A and MR1B transcripts are widely expressed. Despite the widespread expression of MR1A and MR1B, only MR1A can present mycobacterial antigen to MAIT cells. Coexpression of MR1B with MR1A serves to decrease MAIT cell activation following bacterial infection. However, expression of MR1B prior to MR1A lowers total MR1A abundance, suggesting competition between MR1A and MR1B for either ligands or chaperones required for folding and/or trafficking. Finally, we evaluated CD4/CD8 double positive thymocytes expressing surface MR1. Relative *MR1A/MR1B* expression in MR1-expressing thymocytes is associated with their prevalence.

Our results suggest alternative splicing of MR1 represents a means of regulating MAIT activation in response to microbial ligand.

**Funding:** This work was supported by NIH T32HL083808 (EK, GAN, EM), VA Merit Award I01CX001562 (MJH), NIH R01AI29976 (MJH), NIH R01AI048090 (DML), NIH R21AI124225-01A1 (FT) and VA Merit Award I01BX000533 (DML). The contents do not represent the views of the U.S. Department of Veterans Affairs or the United States Government.

## Introduction

MR1-restricted T (MAIT) cells are a highly abundant subset of CD8+ T cells that represent 1-10% of circulating T cells. They are enriched in mucosal sites, including the skin, intestinal mucosa, and lung, and they can serve to sense intracellular microbial infection.(1) MAIT cells produce the effector cytokines IFN *γ* and TNF *α* following encounter with infected cells.(2, 3) MAIT cells respond to a wide array of microbes including, but not limited to, *Mycobacterium tuberculosis* (Mtb), *Escherichia coli (E. coli)*, *Candida albicans (C. albicans)*, *Legionella pneumophila*, and *Streptococcus* species.(2–6) While the prevalence and phenotype of MAIT cells in mice is distinct from that in humans, mice lacking MAIT cells were unable to fully control infection with *F. tularemia*, BCG, Klebsiella, and Legionella longbeachiae, (6–9). In some cases, MAIT cells have been associated with releasing early signals that play a role in the recruitment of activated MHC-II and MHC-I restricted T cells (6, 7). Therefore, MAIT cells represent an important mechanism by which the immune system senses and responds to microbial infection.

MAIT cells recognize microbial antigen processed and presented by the major histocompatibility complex (MHC) Class I related molecule, MR1.(8) MR1, though similar in many ways to canonical Class I molecules, has distinct features that render it uniquely suited to present microbial ligands. *MR1* transcript is expressed in all nucleated cells; however unlike MHC Class I molecules, which are constitutively detected on the cell surface, MR1 resides in the endoplasmic reticulum (ER) and late endosomal vesicles.(9, 10) Following infection, MR1 binds microbial ligand, and this complex is thought to traffic to the cell surface to stimulate MAIT cells.(9, 10) We have previously shown that MR1 mediated antigen presentation is dependent on the vesicular trafficking proteins Syntaxin18 and VAMP4.(11) More recently, we have observed that distinct trafficking pathways exist to present endogenous and exogenous mycobacterial antigen by MR1 to stimulate MAIT cells (11, 12).

While MHC Class I molecules traditionally present peptides to stimulate CD8+ T cell responses, MR1 binds and presents microbial small molecule metabolites to MAIT cells. These antigens were first described as intermediates in the riboflavin synthesis pathway, but recent reports have highlighted the increasing diversity of the MAIT ligand repertoire.(13–15) For example, MR1 also binds and presents antigen from *S. pyogenes*, a bacteria that cannot synthesize riboflavin (4). *In silico* screens showed that MR1 can bind a range of synthetic compounds, including commonly prescribed pharmaceuticals (14). We have recently shown through metabolomics analysis that MAIT stimulatory antigens include ligands distinct from those generated in the riboflavin synthesis pathway (5).

The underlying gene organization of MR1 is distinct from canonical MHC molecules. *MR1* is non polymorphic, highly conserved across species and individuals, with the transcript ubiquitously expressed(16–18). *MR1* pre-mRNA undergoes alternative splicing to produce multiple isoforms, which have been demonstrated at the transcript level to be expressed in human tissues and cell lines (19). The structure of MR1 is similar to that of MHC Class I molecules, with *α*1 and *α*2 domains that bind ligand, an *α*3 domain that interacts with *β*2-microglobulin, and a transmembrane domain for surface expression.(19, 20) The full length isoform, MR1A, contains all encoded exons and can stimulate MAIT cells. The shorter isoform MR1B lacks the *α*3 domain but does encode the ligand binding and transmembrane domains. The function of MR1B is not well characterized. Overexpression of MR1B in a fibrosarcoma model suggested a functional role for MR1B in stimulating MAIT cells following infection with E*. coli* (21). MR1C is a putative soluble isoform, lacking both the *α*3 and transmembrane domains, however, there are no reports on its expression or function.

Here, we sought to determine the role of MR1 isoforms in the presentation of microbial ligand to MAIT cells. We show *MR1A and MR1B* transcripts are detectable across human tissues, with considerable variation in isoform expression among donors and tissues. We developed a lung epithelial cell line deficient in *MR1* and utilized this system to show that MR1B can antagonize MR1A in the presentation and/or processing of mycobacterial antigen(s). While MR1A is observed in the ER and vesicular compartments, MR1B appears to reside primarily in intracellular vesicles. Finally, we show that surface expression of MR1A on CD4+CD8+ thymocytes is associated with relative abundance of these cells Taken together, our results indicate a novel mechanism by which MAIT cell activation is potentially regulated to target immune responses against microbial infection.

## Materials and Methods

### Human subjects

All samples were collected, and all experiments were conducted under protocols approved by the institutional review board at Oregon Health and Science University. PBMCs were obtained by apheresis from healthy adult donors with informed consent. De-identified lungs, upper airway, or small intestine were obtained from the Pacific Northwest Transplant Bank (PNTB). Our exclusion criteria included significant tobacco smoking history (>1 pack-year), drowning, crushing chest injuries, lobar pneumonia, and HIV/HBC/HCV infection. Deidentified thymuses were obtained from children undergoing cardiac surgery at Oregon Health and Science University Doernbecher Children’s Hospital. The majority of children were no less than 4 mo. and no more than 4 years old. As the thymuses were obtained as deidentified medical waste under an exempt IRB protocol, no other information is available on the status of the donors.

### Reagents and antibodies

Doxycycline (Sigma-Aldrich. St. Louis, Missouri) was resuspended to 2mg/mL in sterile water and was used at 2 *µ*g/mL. 6-formylpterin (Schirck’s Laboratories, Bauma, Switzerland) was resuspended to 3mg/mL in 0.01M NaOH. Control vehicle used was 0.01M NaOH.

### Microorganisms and preparation of APCs

*M. smegmatis* and *M. bovis* Bacillus Calmette-Guérin (BCG) were utilized from frozen glycerol stock. For *M.smegmatis* infection, indicated cells were infected for 1h in a 96 well ELISPOT MSHA nitrocellulose plate. For *M.* bovis BCG infection, cells were plated for four hours in a 6 well tissue culture plate and infected overnight at an MOI of 15. Subsequently, *M. bovis* BCG infected cells were harvested, washed, counted, and added to the ELISPOT assay at the indicated amount of antigen presenting cells. To generate *M.smegmatis* supernatant, *M. smegmatis* was cultured with shaking for 24 hours then pelleted. The supernatant was passed through a 0.22 *µ*m filter to remove any bacteria. The supernatant was aliquoted and stored at −80 °C and utilized at the indicated volume in an ELISPOT assay.

### Cell lines

BEAS-2B and A549 cell lines was obtained from ATCC. A549_MR1KO were generated as described previously (22). All cell lines were cultured in DMEM (Gibco) supplemented with L-glutamine and 10% heat inactivated FBS. All cell lines were confirmed to be mycoplasma free.

### Generation of lentivirus particles

Lentivirus production was performed as described previously (23). Briefly, lentiviruses were produced by co-transfection of HEK 293T cells with the lentiviral vectors that contain our gene of interest (pCDH-sgRNA or pCW-Cas9) and the packaging plasmids (psPAX2 and pMD2.G). Transfection was performed with Lipofectamine 3000 (Thermo Fisher) according to manufacturer’s instructions. Cells were cultured in DMEM supplemented with 10%FBS and the growth medium was replaced after 6 hours. 48 hours after transfection, lentivirus-containing supernatants were harvested, centrifuge for 5min at 1250 rpm and filtered through a 0.45*µ*m filter.

### Generation of a Beas2B MR1 knockout cell line

CRISPR/Cas9-mediated genome-editing for MR1 was performed as described previously (23). The wildtype Beas2B cells were seeded in a 6 well plate to 70% confluency and transduced by spinoculation with a lentivirus encoding the CRISPR/Cas9 gene with guide RNA specifically targeted to MR1 that had been previously used to generate an A549 MR1 knockout cell line (22). Following this, limiting dilution was performed on transduced cells, with 1 cell per well in 96 well plates. Cells were grown in DMEM media containing 10% fetal bovine serum and 1% gentamicin antibiotic. Clones were utilized as antigen presenting cells in an ELISPOT assay following incubation with *M.smegmatis* supernatant and a MAIT clone as described below. Positive clones were expanded in a 12 well plate and tested again for knockout of MR1 by ELISPOT. Following this, functionally knocked out clones were incubated with 100 *µ*M 6FP versus an equivalent volume of 0.01M NaOH. Surface staining of MR1 was performed as described below and MR1 surface expression was assessed by flow cytometry on a BD Fortessa. Three clones were validated as knocked out for MR1 protein expression, as well as functionally unable to stimulate MAIT clones, and cryopreserved in 90% fetal bovine serum/10% DMSO. One clone was selected and utilized for all future experiments (Beas2B_MR1KO).

### Generation of stably transduced Beas2B cell lines

Lentivirus production was performed as described previously (24). Briefly, lentiviruses were produced by co-transfection of HEK 293T cells with the lentiviral vectors that contain our gene of interest (pCDH-sgRNA or pCW-Cas9) and the packaging plasmids (psPAX2 and pMD2.G). The Beas2B_MR1KO cell line was transduced with a lentivirus expressing pCI_Tet_MR1AGFP by spinoculation as described by Tafesse et al (24). Transduced cells were exposed to doxycycline overnight to promote expression of MR1AGFP and subsequently sorted on an InFlux cell sorter based on GFP expression. Cells were subsequently cultured, validated for MR1AGFP expression by flow cytometry following administration of doxycycline, and utilized for further experiments.

### Monocyte-derived DCs

PBMCs obtained by apheresis were resuspended in 2% human serum in RPMI and were allowed to adhere to a T-75 flask at 37 °C for 1 h. After gentle washing twice with PBS, nonadherent cells were removed and 10% human serum in RPMI containing 30 ng ml^−1^ of IL-4 (Immunex) and 30 ng ml^−1^ of granulocyte–macrophage colony-stimulating factor (Immunex) was added to the adherent cells. The cells were X-rayed with 3,000 cGray using X-RAD320 (Precision X-Ray Inc.) to prevent cell division. After 5 days, cells were harvested with cell-dissociation medium (Sigma-Aldrich, Gillingham, UK) and used as APCs in assays.

### Human tissue sources of antigen presenting cells

PBMCs were isolated from the peripheral blood of healthy donors using Ficoll-Paque gradients.

Lung and small intestine single cell suspensions are prepared from recently deceased donor tissue not suitable for transplant from the Pacific Northwest Transplant bank. Small cubes of lung parenchyma, devoid of airway and lymph nodes, or of duodenal lamina propria, were cut into a cold buffer of HBSS (Gibco) media supplemented with HEPES (Gibco) and PSF antibiotic (Sigma). Tissue was then digested for 30 minutes at 37°C in a DMEM buffer (Gibco) supplemented with PSF antibiotics (Sigma), elastase (15 *µ*g/mL, Worthington), trypsin I (1.5 *µ*g/mL, Sigma), DNase I (45 *µ*g/mL, Roche). The subsequent suspension was further dissociated using a GentleMACS dissociator (Miltenyi). The single cell suspension was then diluted 1:1 with a buffer of HBSS (Gibco) media supplemented with 2% heat-inactivated fetal bovine serum (Gemini Bio Products), HEPES (Gibco) and PSF antibiotic (Sigma) to dilute homogenate and neutralize digest enzymes. This cell suspension is passed through successive filters in this order: metal mesh sieve filter (size 40 then 60, Sigma), and nylon cell strainer (100um then 40 *µ*m, BD Falcon). The resulting cell suspension is washed in RPMI supplemented with 10% heat inactivated human serum and used for experiments or cryo-preserved in heat-inactivated fetal bovine serum with 10% DMSO. Large airway epithelial cells were generated as described previously from the upper airway of human donors and cryopreserved in heat-inactivated fetal bovine serum with 10% DMSO.(2)

Thymocytes: Thymus tissue was cut into 3-mm^3^ pieces. Each piece was ground in a GentleMACS dissociator with 1 ml of DMEM plus 10% FBS to form a single cell suspension. The suspension was cryopreserved at 2-3×10^8^ cells/ml in a 90% FBS/10% DMSO freezing solution with a post-thaw viability of approximately 50%. ^22^

### Expansion of T-cell clones

T-cell clones were cultured in the presence of X-rayed (3,000 cGray using X-RAD320, Precision X-Ray Inc.) allogeneic PBMCs, X-rayed allogeneic LCL (6,000 cGray) and anti-CD3 monoclonal antibody (20 ng ml^−1^; Orthoclone OKT3, eBioscience) in RPMI 1640 media with 10% human serum in a T-25 upright flask in a total volume of 30 ml. The cultures were supplemented with IL-2 on days 1, 4, 7 and 10 of culture. The cell cultures were washed on day 5 to remove soluble anti-CD3 monoclonal antibodies.

### IFN-***γ*** ELISPOT

A MSHA S4510 96 well nitrocellulose-backed plate (Millipore, bought via Fisher Scientific) was coated overnight at 4 °C with 10 μg ml^−1^solution of anti-IFN-γ monoclonal antibody (Mabtech clone 1-D1K) in a buffer solution of 0.1 M Na2CO3, 0.1 M NaHCO3, pH=9.6). Then, the plate was washed three times with sterile PBS and blocked for 1 h at room temperature with RPMI 1640 media containing 10% heat-inactivated HS pool. Then, the APCs and T cells were prepared as described above and co-incubated overnight. Antigen presenting cells and antigen used are noted in the relevant figure. T-cell clones were added at 5 × 10^3^ per well. The plate was incubated overnight at 37 °C and then washed six times with PBS containing 0.05% Tween. The plate was then developed as previously described and analyzed using an AID ELISPOT reader.(5)

### Construction of MR1B_RFP, MR1C_RFP and transfections

We utilized our previously described pCI construct for all transient transfections (10). A gene encoding MR1B_RFP or MR1C_RFP was ligated into this construct using EcoRI and KpnI, to create pCI_MR1B_RFP or pCI_MR1C_RFP. Confirmatory sequencing was performed. Restriction enzymes and ligation kit were obtained from New England Biolabs (Ipswich, Massachusetts). PCR and gel purification kits were obtained from Qiagen. Transfection of Beas2B or A549 cells using plasmids was done using an Amaxa Nucleofector, Kit T solution (Lonza, Basel, Switzerland), and programs G-016. Each transfection reaction was done with 1e6 cells and 5 *µ*g of plasmid. For co transfections, equivalent molar amounts of each plasmid were used for a total of 5 ug of plasmid. A pCI empty vector was used as a control for all transfections. Confirmation of transfection was performed using flow cytometry on a BD Fortessa to detect either GFP (pCI_MR1AGFP) or RFP (pCI_MR1BRFP or pCI_MR1CRFP). All analyses were performed using FlowJo software (TreeStar).

### RNA isolation, cDNA synthesis and qPCR analysis

Total RNA was isolated using TRIZOL (Life Technologies) and phenol chloroform extraction followed by an RNAeasy Kit (Qiagen). cDNA was synthesized using a High Capacity cDNA Reverse Transcription Kit (ThermoFisher Scientific). qPCR was performed using SYBRGreen Power Master Mix (ThermoFisher Scientific) on a Step One Plus Real-Time PCR System (Applied Biosystems). Primers were designed to be specific for each gene assay and generated by the DNA services core and IDT Technologies. The absolute quantification method was used using by generating a standard curve with a plasmid specific for each gene assayed. Primer sequences are noted in Table 3.

**Table 1:**
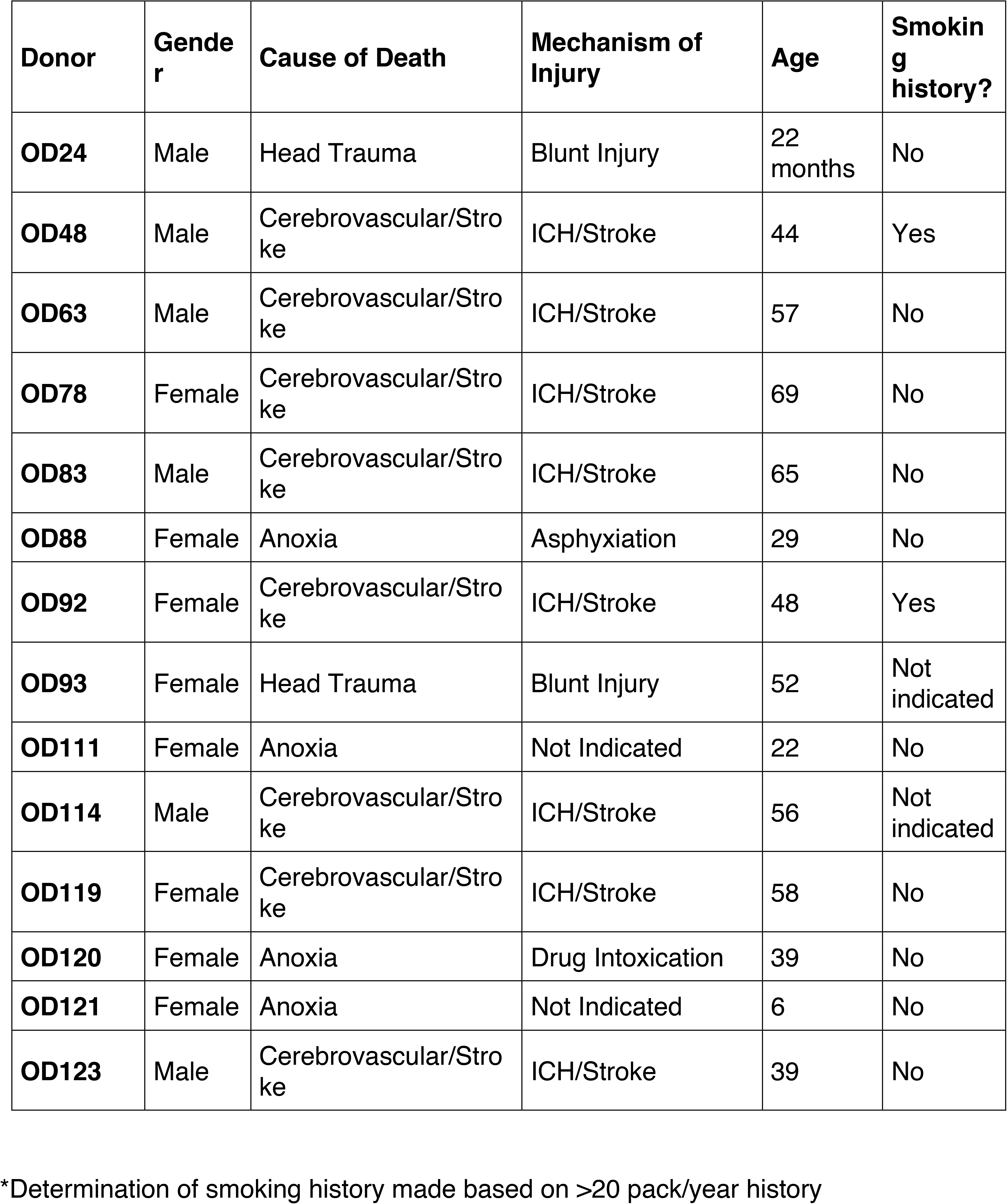
Table of airway, lung, and small intestine lamina propria donors.

| **Donor** | **Gender** | **Cause of Death** | **Mechanism of Injury** | **Age** | **Smoking history?** |
| --- | --- | --- | --- | --- | --- |
| **OD24** | Male | Head Trauma | Blunt Injury | 22 months | No |
| **OD48** | Male | Cerebrovascular/Stroke | ICH/Stroke | 44 | Yes |
| **OD63** | Male | Cerebrovascular/Stroke | ICH/Stroke | 57 | No |
| **OD78** | Female | Cerebrovascular/Stroke | ICH/Stroke | 69 | No |
| **OD83** | Male | Cerebrovascular/Stroke | ICH/Stroke | 65 | No |
| **OD88** | Female | Anoxia | Asphyxiation | 29 | No |
| **OD92** | Female | Cerebrovascular/Stroke | ICH/Stroke | 48 | Yes |
| **OD93** | Female | Head Trauma | Blunt Injury | 52 | Not indicated |
| **OD111** | Female | Anoxia | Not Indicated | 22 | No |
| **OD114** | Male | Cerebrovascular/Stroke | ICH/Stroke | 56 | Not indicated |
| **OD119** | Female | Cerebrovascular/Stroke | ICH/Stroke | 58 | No |
| **OD120** | Female | Anoxia | Drug Intoxication | 39 | No |
| **OD121** | Female | Anoxia | Not Indicated | 6 | No |
| **OD123** | Male | Cerebrovascular/Stroke | ICH/Stroke | 39 | No |

\*Determination of smoking history made based on >20 pack/year history

**Table 2:** Antibodies used in flow cytometry surface staining.

| **<u>Antibody Name</u>** | **<u>Fluorophore</u>** | **<u>Manufacturer</u>** | **<u>Clone</u>** |
| --- | --- | --- | --- |
| CD4 | PE Cy7 | BioLegend | RPA-T4 |
| CD326 | APC | BioLegend | 9C4 |
| CD14 | FITC | Biolegend | HCD14 |
| CD20 | FITC | BioLegend | 2H7 |
| CD45 | FITC | Biolegend | 30-F11 |
| Live-Dead | Aqua | ThermoFisher |  |
| MR1/5-OP-RU tetramer | PE | NIH Tetramer<br>Core (Emory) |  |
| MR1/6FP tetramer | PE | NIH Tetramer<br>Core (Emory) |  |

**Table 3:**
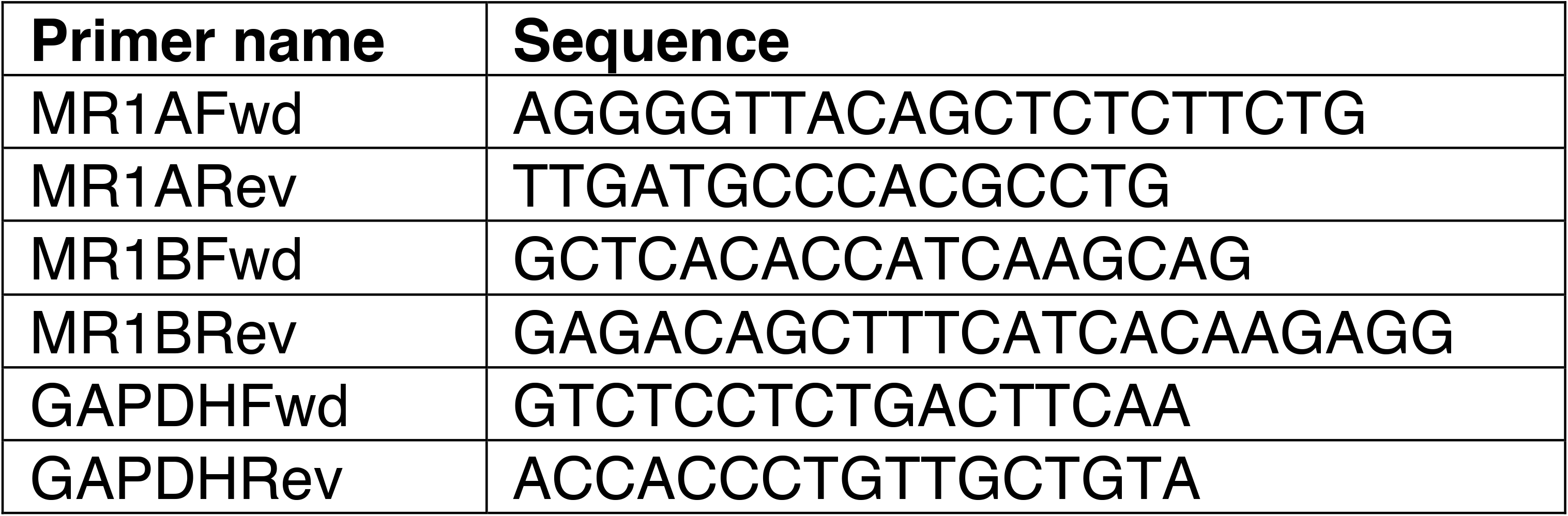
Primer pairs used for qRT-PCR.

**Table 4:** Frequencies of MR1+ and MAIT populations in the thymus.

| **Donor** | **Prevalence of MR1 expressing thymocytes (% of DP thymocytes)** | **MR1 protein expression (gMFI MR1+ cells)** | **Prevalence of MAITs (% of 5-OP-RU Tetramer+ of Live CD3)** |
| --- | --- | --- | --- |
| 90 | 5.33 | 1529 | 0.048 |
| 97 | 6.77 | 1466 | 0.200 |
| 99 | 8.00 | 1337 | 0.150 |
| 106 | 2.67 | 788 | 0.020 |
| 107 | 2.00 | 816 | 0.020 |
| 110 | 0.50 | 1060 | 0.058 |
| 118 | 0.75 | 873 | 0.018 |
| 121 | 0.10 | 796 | 0.130 |
| 152 | 1.87 | 644 | 0.130 |

### MR1 surface stabilization

BEAS-2B:MR1_KO cells, either untransfected, or transfected with pCI empty vector, pCI_MR1AGFP, and/or pCI_MR1BRFP, were plated into 6 well tissue culture plates overnight. The next day, the cells were treated with 100 *µ*M 6-FP versus an equivalent volume of 0.01M NaOH. After 16 hours, the cells were harvested on ice and split into two groups for primary staining and isotype control staining. Primary staining was done with an antibody against MR1 (Clone 26.5, gift from Ted Hansen, biotinylated by Biolegend) at 1:100 for 40 min on ice in the presence of 2% human serum, 2% goat serum, and 0.5% FBS. Biotinylated mouse anti IgG2A (Biolegend) served as the isotype control. After washing, streptavidin-Alexa 647 (ThermoFisher Scientific) was added for 40 min on ice. Cells were washed, fixed in 1% PFA and analyzed by flow cytometry on a BD Fortessa, or BD Symphony and data were analyzed on FlowJo (TreeStar).

### Fluorescence microscopy

BEAS-2B:MR1_KO cells were transfected with MR1A_GFP or MR1B_RFP and plated into 1.5mm glass bottom chamber slides (Nunc, ThermoFisher Scientific) and incubated at 37°C and 5% CO_2_. After 48h, live images were acquired on a high-resolution wide field CoreDV system (Applied Precision, Pittsburgh, Pennsylvania) with a Nikon Coolsnap ES2 HQ. For 6FP surface stabilization assay, cells were transfected as above, and administered 50 uM 6FP or an equivalent volume of 0.01 M NaOH for 16h. Cells were fixed for 20 minutes in 4% PFA and stained with DAPI (ThermoFisher Scientific) to visualize nucleus before acquisition on a wide field CoreDV system. Each image was acquired as Z-stacks in a 1024×1024 format with a 60x objective (NA 1.42).

### Dynamics of MR1A and MR1B expression

To assess the temporal dynamics of MR1A and MR1B expression, pCI_MR1BRFP or pCI_MR1CRFP was transfected into Beas2B_MR1KO cells overexpressing a Tet_MR1AGFP lentivirus either 24h prior to or 16 hours post induction of MR1A expression. Flow cytometry was performed as described earlier to measure MR1AGFP expression or MR1BRFP expression using a BD Symphony and data were analyzed using FlowJo software. For functional analyses, cells were harvested on ice, infected 1h with a titrating dose of M. smegmatis, and utilized as antigen presenting cells (5X10^3/well) in an ELISPOT assay following incubation with an MAIT clone (5X10^3/well). The ELISPOT was set up and developed as described above.

### Flow cytometry staining and FACS sorting

Lung and small intestine cell suspensions generated as described earlier were thawed, washed with PBS twice and blocked in FACS buffer. 2e06 cells were stained with antibodies against surface CD45, CD326 (EPCam), CD14, and CD20, for 30 minutes on ice in the dark, as well as Live/Dead Fix Aqua for viability. Cells were harvested on ice, washed in PBS, resuspended in FACS buffer, and sorted on a BD InFlux sorter for Live, CD45-CD14-, and CD20-CD326+ cells in order to isolate CD326+ epithelial cells. Cells were sorted into TRIZOL for immediate RNA isolation and subsequent quantitative real time PCR analysis.

Sorting of MR1+ DP thymocytes: Thymocytes were thawed and blocked with FACS buffer, and 2e06 cells were stained with antibodies for the surface markers CD3, CD4, CD8, and MR1-biotin (clone 26.5), or the isotype control (IgG2a-biotin) for 30 minutes on ice in the dark. Cells were washed twice with cold PBS and stained with streptavidin-PE (1:2000 dilution) for 15 minutes in the dark. Cells were washed twice and resuspended in FACS buffer and sorted on a BD InFlux Sorter for Live, CD3+ CD4+CD8+ MR1 expressing thymocytes, which was confirmed based on an isotype control for MR1. Cells were sorted into TRIZOL for immediate RNA isolation and subsequent quantitative real time PCR analysis.

For quantifying the frequency of MAIT thymocytes, thymocytes were thawed, blocked with FACS buffer, and 2e06 cells were incubated with the MR1/5-OP-RU tetramer or the MR1/6FP tetramer (1:500 dilution, NIH Tetramer Core facility), for 45 minutes at RT in the dark. Following this, antibodies to the surface markers CD3, CD4, and CD8 and Live/Dead Fix Aqua viability stain were added at manufacturer recommended concentrations. Cells were incubated at RT in the dark for 15 minutes, washed twice with PBS, and fixed with 1% paraformaldehyde. Acquisition was performed on a BD Symphony flow cytometer and data were analyzed using FlowJo software (TreeStar).

### GTEx analysis

Figure 1 depicts the approximate ratio of exon 4 inclusions to skips in *MR1* across normal tissue RNA-seq samples comprising the GTEx project.(25) To generate this figure, we first used Snaptron to query GTEx RNA-seq samples for numbers of reads spanning each of three exon-exon junctions, whose *hg38* coordinates are a) chr1:181050287-181052234, b) chr1:181052511-181053572, and c) chr1:181050287-181053572. Junctions a) and b) point to inclusion of exon 4, while junction c) points to exclusion of exon 4. For a given sample, we averaged the numbers of reads spanning junctions a) and b) and divided the result by the number of reads spanning junction c) to estimate the ratio of exon 4 inclusion to exclusion.(26) We subsequently used *Mathematica* 10.4 to plot results across samples. Scripts to reproduce Figure 1 are available at https://github.com/nellore/mr1.

**Figure 1:**
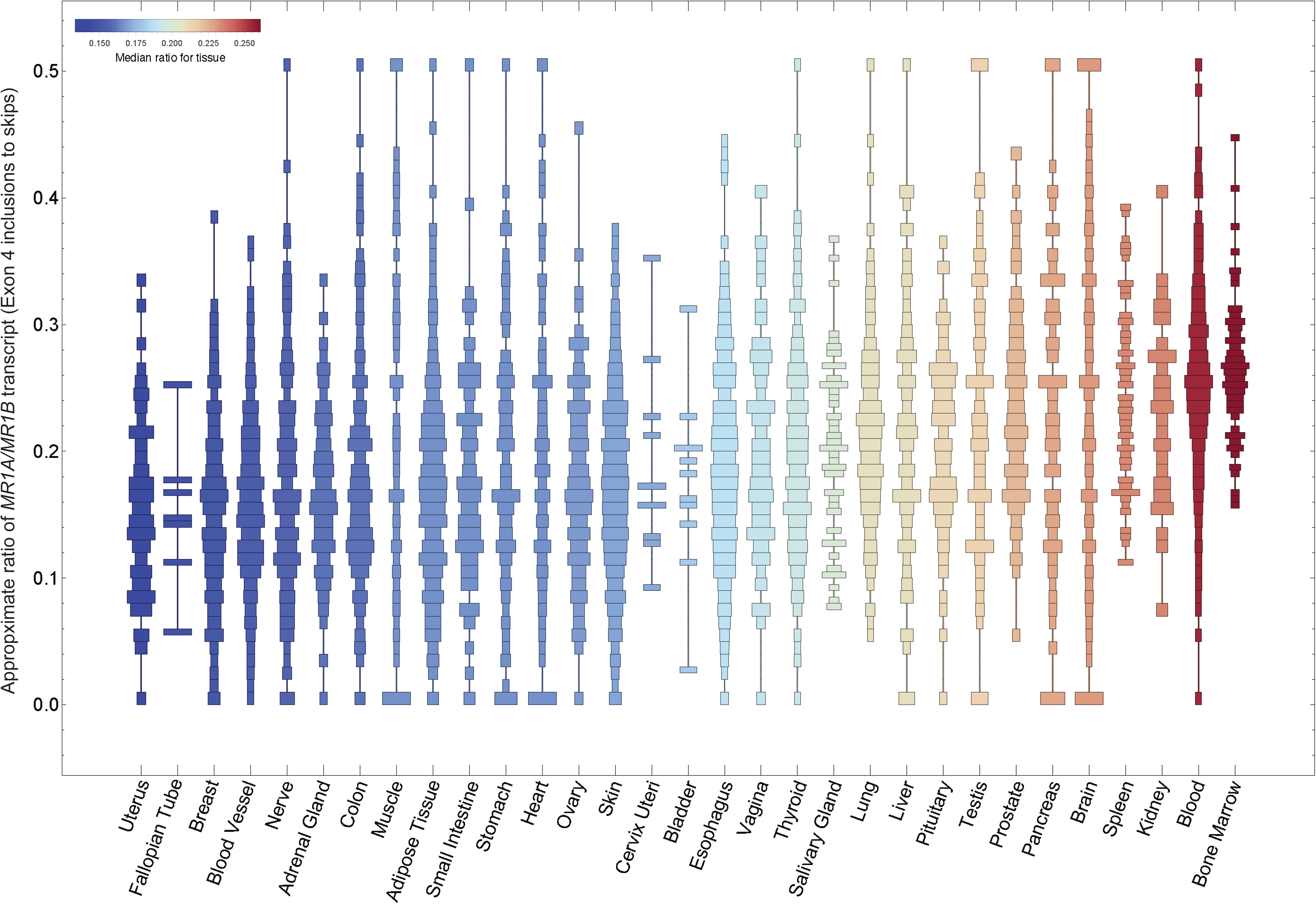
Distribution of relative *MR1A/MR1B* transcript across the GTex dataset. Snaptron was used to query human transcriptome data from the publicly available GTEx dataset of non-malignant human tissues. Relative *MR1A/MR1B* mRNA expression was measured by quantifying junctional inclusion ratios of exon 3 inclusions (*MR1A*) to exon 3 skips (*MR1B*). We depict histograms of the ratio of *MR1A/MR1B* transcript expression in order of increasing mean expression.

## Data analysis

Data were analyzed and plotted using Prism 7 GraphPad Software (La Jolla, California). Statistical significance was determined using unpaired Student’s two-tailed *t*-test, unless otherwise indicated. For comparison of DP MR1+ thymocyte gene expression with frequency of DP thymocytes or of MAIT thymocytes, Pearson’s correlation calculation was performed. For comparisons of functionality between transfected antigen presenting cells, linear regression analysis was performed and significant differences between slopes and y-intercepts were measured and reported. Error bars in the figures indicate the standard deviation, standard error of the mean, or the data set range as indicated in each figure legend. *P* values < 0.05 were considered significant (\**P* < 0.05; ** *P* < 0.01; *** *P* < 0.001)

## Results

### MR1A and MR1B are ubiquitously expressed

To study expression of the MR1 splice variants, we used Snaptron, a tool for exploring exon-exon junction expression across thousands of publicly available RNA sequencing (RNA-seq) samples.(26, 27) We looked at exon-exon junctions corresponding to inclusion or exclusion of exon 4 to distinguish *MR1A* from *MR1B*, using nearly 10,000 RNA-seq samples from 31 non-diseased tissue sites in over 500 deceased individuals from the Genotype Tissue Expression (GTEx) project (see Materials and Methods).(25) Here, we show that both *MR1A* and *MR1B* are expressed across human tissues, as seen in prior, non-quantitative studies (Figure 1).(16, 19, 28) The ratio of *MR1A* to *MR1B* varied across tissues, with higher relative *MR1A* transcript observed in blood, bone marrow, liver, and lung, and lower relative *MR1A* observed in uterine cervix, breast, small intestine, and colon. Interestingly, we observed that in all the tissues queried, the *MR1A/MR1B* ratio was consistently less than 0.5, suggesting that all the tissues express higher relative *MR1B* than *MR1A.* As these data are from the total mRNA for a given tissue, this analysis does not take into account diversity of individual cell types that comprise each tissue.

### Quantitative RT-PCR analysis reveals differing levels of MR1 splice variants in antigen presenting cells

To explore the relative MR1 splice variant expression specifically in antigen presenting cells (APCs), we utilized quantitative real-time PCR (qRT-PCR) following isolation of APCs from human blood and lung tissue. We generated a standard curve for each splice variant by using plasmids designed to be specific to each isoform (Figure 2A). In this manner, we were able to quantify absolute and relative amounts of the MR1 splice variants. From PBMC, we generated monocytes which were then differentiated to generate dendritic cells or macrophages.(29) We observed that in both dendritic cells (DCs) and macrophages, both *MR1A* and *MR1B* transcripts were readily detected (Figure 2B).

**Figure 2:**
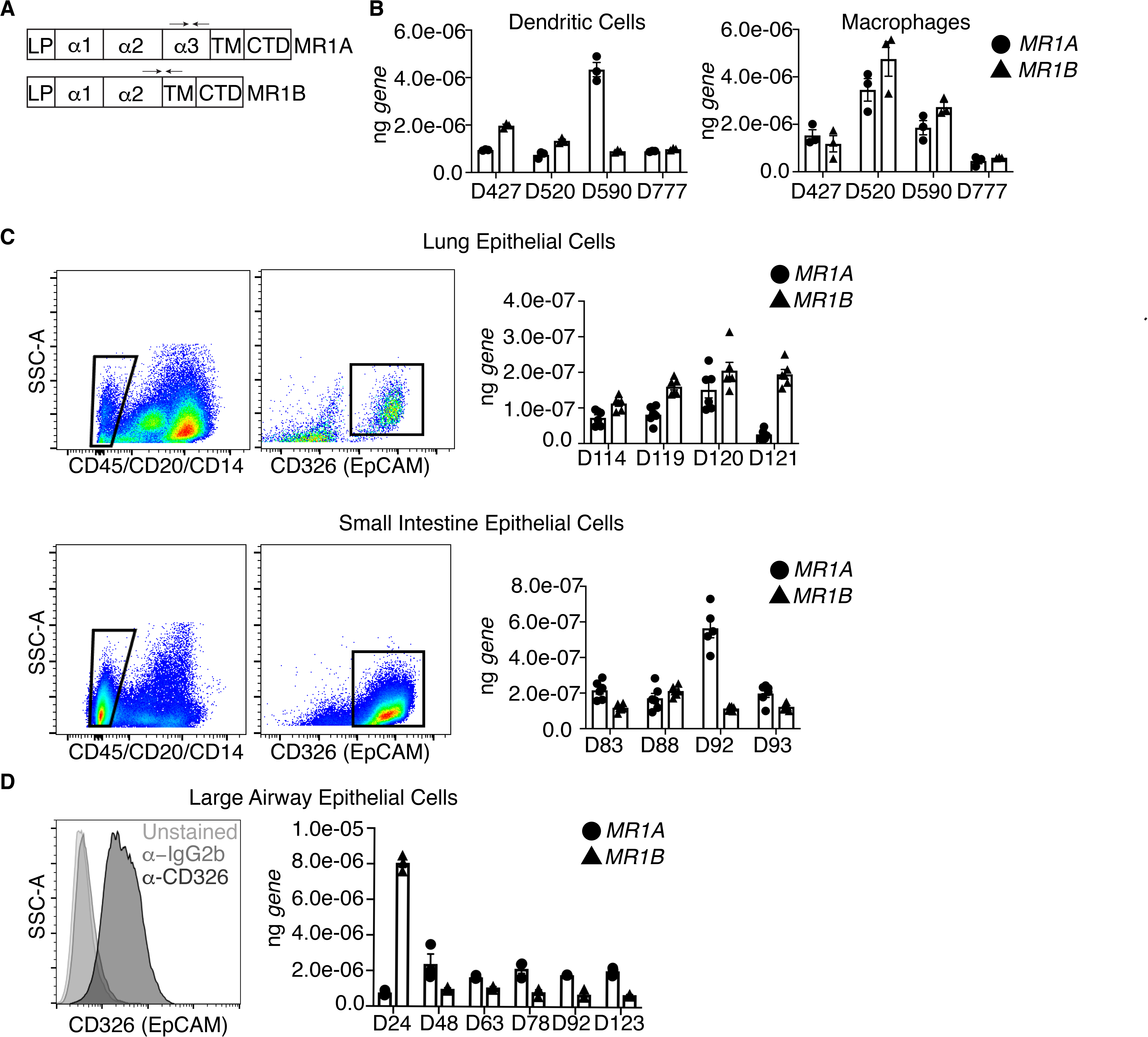
MR1A and MR1B mRNA are detectable in antigen presenting cells isolated from human tissues. Quantitative real-time PCR (qRT-PCR) primers were designed to specifically amplify either *MR1A* or *MR1B* transcript. (A). In brief: RNA was isolated from 5e04 cells and qRT-PCR was performed to quantify the absolute amounts of *MR1A* or *MR1B* mRNA as measured by a standard curve designed to be specific for each isoform. (B) qRT-PCR performed on monocyte-derived macrophages and monocyte-derived dendritic cells generated from human PBMC (C) Human lung parenchyma and small intestine lamina propria epithelial cells were stained with antibodies against CD45, CD20, CD14 and CD326 (EpCAM). CD45-CD20-CD14-CD326+ cells were sorted for mRNA isolation and cDNA synthesis. qRT-PCR was performed as described (D) Human Large Airway Epithelial Cells (LAEC) were isolated and cultured from the upper airway and stained with an antibody to either isotype (IgG2b) or CD326 (EpCAM) to verify surface expression of EpCAM. mRNA was isolated and qRT-PCR performed as described. Individual replicates are shown and error bars represent mean and standard error.

To determine the expression of MR1 splice variants in primary human epithelial cells, we generated single cell suspensions from lung parenchyma and small intestine and subsequently sorted epithelial cells by EpCam (CD326) staining (Figure 2C). Using qRT-PCR we found that expression of *MR1A* was again similar to *MR1B*. Interestingly, both lung and small intestine cells expressed nearly 2-log lower levels of *MR1* splice variants than cells generated from PBMC.

As primary large airway epithelial cells (LAEC) are capable of presenting mycobacterial antigen to stimulate MAIT cells, we prepared LAEC from human airway tissue, and confirmed that these cells expressed for EpCam (Figure 2D).(2) LAEC isolated from 5/6 donors expressed relatively more *MR1A* mRNA compared to *MR1B.* Overall transcript expression of these LAEC MR1 splice variants was similar to levels observed in APCs generated from PBMC. *MR1A* mRNA levels were more consistent across donors, ranging from 1.0E-06 ng to 2.1E-06 ng, while *MR1B* transcript ranged from 2E-07 ng to 4E-06 ng. Of note, LAEC from one donor, D24, expressed considerably higher (4-fold) expression of *MR1B* than *MR1A*, relative to the LAEC prepared from the remaining donors which expressed 2-5-fold higher *MR1A* relative to *MR1B.* Taken together, these results confirm that both *MR1A* and *MR1B* transcript are expressed, but highlights heterogeneity both within tissues and across donors with regard to the expression of *MR1* splice variants.

### MR1B inhibits T cell activation by MR1A

We next sought to assess the function of the MR1 isoforms in the absence of endogenous MR1. Here, a previously described CRISPR/Cas9 method utilized in A549 cells was used to generate a bronchial epithelial cell line (Beas2B) that lacked the gene for *MR1* (Beas2B:MR1_KO).(22) Following limiting dilution, we used MR1 surface stabilization following treatment with 6FP to identify Beas2B:MR1 KO cells (Figure 3A).(30) Functionally, we found that these clones could not stimulate an MR1-restricted clone in response to *M. smegmatis* infected APC (Figure 3B). In contrast, stimulation of both HLA-B45 and HLA-E restricted clones was not affected (Figure 3B).

**Figure 3:**
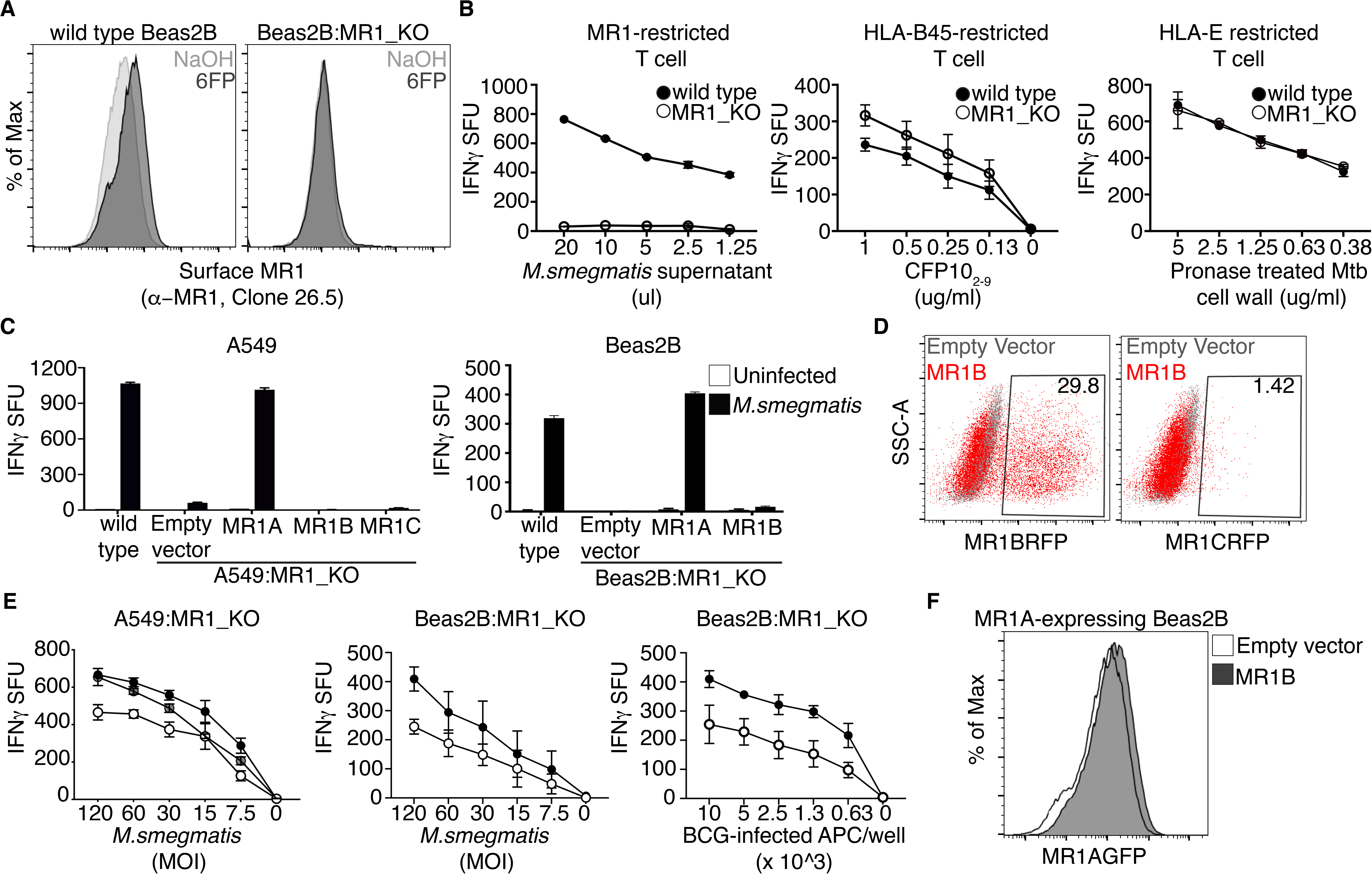
MR1B inhibits T cell activation by MR1A. Wild type Beas2B cells were transduced with a lentivirus targeting *MR1* using CRISPR/Cas9 gene editing to generate a Beas2B MR1 knockout cell line (Beas2B_MR1KO). (A) Surface expression of MR1A assessed using flow cytometry of antibody staining to MR1A (*α*-MR1, 26.5) following exposure of wt Beas2B and Beas2B_MR1KO cells to 6FP overnight. (B) wt Beas2B and Beas2B_MR1KO cells were treated with either *M.smegmatis* supernatant (left), pronase treated Mtb cell wall (middle), or CFP102-9 and (right) utilized to stimulate either a MR1-restricted T cell clone (left), an HLA-E restricted T cell clone (middle), or a HLA-B45 restricted T cell clone (right). IFN-*γ* production is measured by ELISpot and reported as IFN-*γ* spot forming units/5000 T cells (IFN-*γ* SFU). Error bars represent mean and standard error from duplicate wells. (C) A549_MR1KO and Beas2B_MR1KO cells were transfected with plasmids encoding either MR1AGFP, MR1BRFP or MR1CRFP, or a pCI empty vector. Cells were infected overnight with *M. smegmatis* at a multiplicity of infection (MOI) of 3 and utilized as antigen presenting cells to stimulate MAIT production of IFN-*γ* in an ELISpot as described in (B). (D) Beas2B_MR1KO cells were transfected as described and MR1B or MR1C expression was measured by detection of total RFP by flow cytometry. (E) A549_MR1KO or Beas2B_MR1KO cells were cotransfected with plasmids encoding MR1AGFP and the pCI empty vector, MR1BRFP, or MR1CRFP. Cells were infected for 1h with *M. smegmatis* and used as in (B). Data are pooled from at least 3 independent experiments with duplicate wells and error bars represent mean and standard error. (F) Beas2B_MR1KO cells were cotransfected as described in (E), infected with *M. bovis* BCG overnight, and used to stimulate IFN-*γ* production by MAIT cells. Data are pooled from at least 2 independent experiments with duplicate wells, and error bars represent mean and standard error. (G) Beas2B overexpressing MR1AGFP were transiently transfected with either the pCI empty vector or *pCI_MR1BRFP*. Flow cytometric analysis was performed to quantify total MR1A expression, as measured by detection of GFP, following transfection.

We next generated expression vectors containing the genes for *MR1A*, *MR1B,* or *MR1C* under the control of the CMV promoter. To distinguish MR1A from MR1B or MR1C following transfection, we tagged MR1A with GFP and MR1B and MR1C with RFP (pCI:MR1AGFP, pCI:MR1BRFP, pCI:MR1CRFP). We then transfected these plasmids individually into Beas2B:MR1_KO and A549:MR1_KO cells and infected the cells with *M. smegmatis* overnight in order to stimulate MAIT cells (Figure 3C). Flow cytometry was used to confirm protein expression. While MR1C protein was detectable by flow cytometry, it was expressed at markedly lower levels than MR1B (Figure 3D). Compared to wild type Beas2B cells, Beas2BMR1_KO cells were incapable of stimulating MAIT cell production of IFN-*γ*. Expression of pCI:MR1AGFP reconstituted MAIT cell release of IFN*γ* in Beas2B:MR1_KO and A549:MR1_KO cells, while expression of the pCI empty vector, pCI:MR1BRFP or pCI:MR1CRFP did not result in MAIT production of IFN-*γ* Figure 3C).

As we found that neither MR1B nor MR1C could present antigen to MAITs, we postulated that these isoforms could work in concert with MR1A to modulate MAIT activation. To test this hypothesis, we coexpressed pCI:MR1AGFP with pCI:MR1BRFP or pCI:MR1CRFP in Beas2B:MR1_KO and A549:MR1_KO cells, and subsequently infected the cells with *M. smegmatis*. Coexpression of MR1A with MR1B resulted in a roughly 40% decrease in MAIT cell activation in both cell lines (Figure 3E). This inhibition was also seen upon infection with *M. bovis* BCG for Beas2B:MR1_KO cells. However, coexpression of MR1C with MR1A did not lead to inhibition of MR1A-mediated presentation of *M.smegmatis-*derived ligand. Co-expression of MR1B or MR1C with MR1A did not result in a decrease in MR1A surface protein abundance as measured by flow cytometry (Figure 3F). Taken together, these results indicate that MR1B may be functioning specifically to inhibit T cell activation by MR1A.

### MR1B resides in the same intracellular compartments as MR1A

To elucidate the subcellular localization of MR1A and MR1B, we individually expressed pCI:MR1AGFP or pCI:MR1BRFP in Beas2B:MR1_KO cells and performed live cell imaging (Figure 4A). MR1A was detectable in both the endoplasmic reticulum as well as intracellular vesicles, as previously described.(9, 11, 12) MR1B, additionally, resided inside these same compartments. When Beas2B:MR1_KO cells coexpressed pCI:MR1AGFP and pCI:MR1BRFP, we observed that MR1A and MR1B resided in the same intracellular compartments (Figure 4B). This suggests that MR1B may be mediating its effects in close proximity to intracellular MR1A. It has been previously established that translocation of MR1A to the cell surface is dependent on the availability of ligands such as 6FP.(11, 30) As MR1A and MR1B reside in the same intracellular compartments, we postulated that MR1B could interfere with the translocation of MR1A to the cell surface, or could directly interfere with the engagement with the MAIT T cell receptor on the cell surface. Beas2B:MR1_KO cells transfected with either pCI:MR1AGFP or pCI:MR1BRFP were exposed to 6FP overnight and imaging was performed to detect cell surface translocation. While we observed cell surface expression of MR1A, MR1B did not traffic to the cell surface upon exposure to exogenous ligand, remaining in intracellular vesicles (Figure 4C). Therefore, these results suggest that MR1B-mediated antagonism is mediated intracellularly, prior to MR1A translocation to the cell surface.

**Figure 4:**
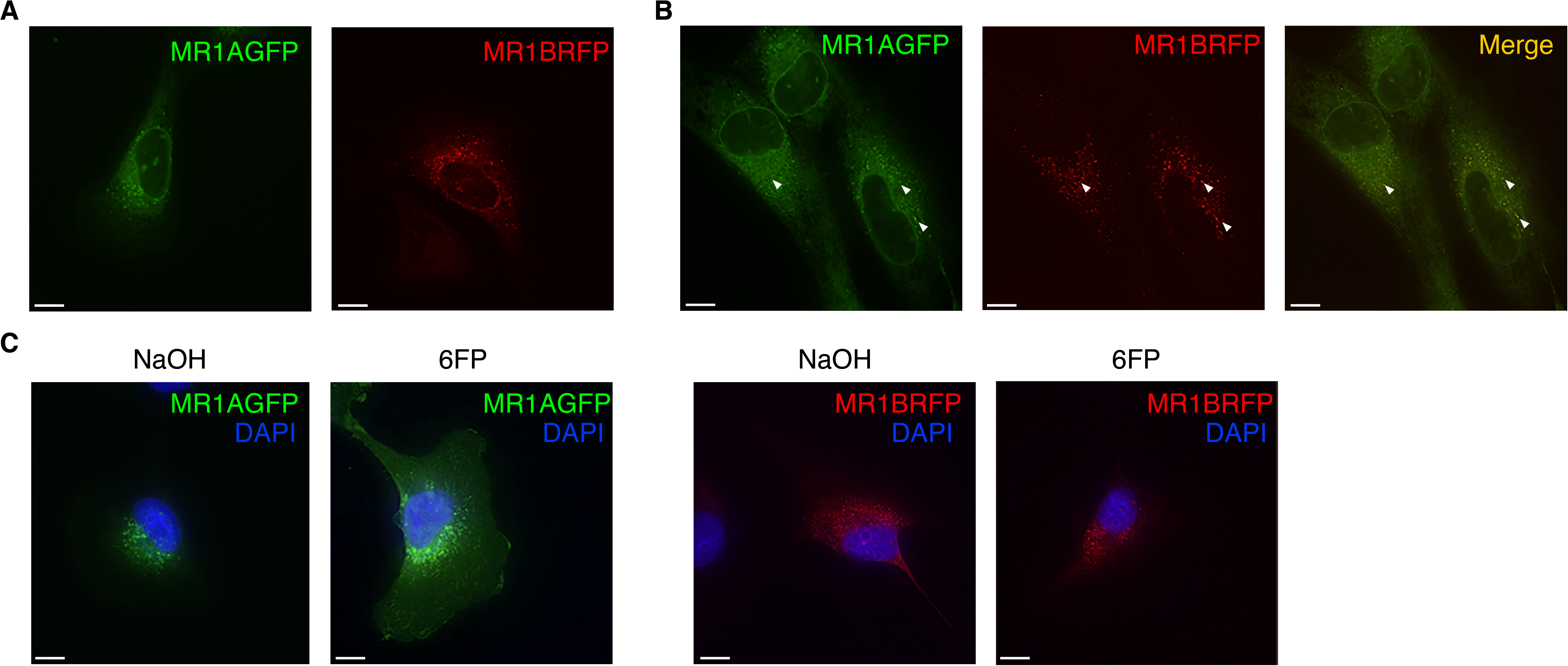
MR1A and MR1B reside in the same intracellular compartments. (A) Beas2B cells were transfected with plasmids encoding either MR1AGFP or MR1BRFP. Live cell imaging was performed using a CoreDV microscope to detect intracellular MR1A (left) or MR1B (right). Scale bars represent 10 *µ*m. (B) MR1A-overexpressing Beas2B cells were transfected with a plasmid encoding MR1BRFP and live cell imaging was performed as described in (A) to determine the subcellular localization of MR1A and MR1B. (left) MR1A alone, (middle) MR1B alone, right (merge). (C) Beas2B cells were transfected with plasmids encoding either MR1AGFP or MR1BRFP, and subsequently treated with 50 *µ*m 6FP or NaOH for 16 hours. Cells were fixed for 20 minutes with 4% PFA and imaging was performed using a CoreDV microscope to determine localization of MR1A or MR1B with or without 6FP. DAPI nuclear stain was used to identify the nucleus. Scale bars represent 10 *µ*m.

### MR1B expression limits abundance of MR1A protein

To further explore the mechanism that MR1B may be inhibiting MR1A abundance or expression, we utilized a doxycycline-inducible MR1A construct. To accomplish this, we stably transduced Beas2B:MR1_KO cells with a lentivirus encoding a doxycycline-inducible MR1AGFP (Beas2B:KO_Tet_MR1AGFP) and sorted the transduced cells based on GFP expression following overnight administration of doxycycline (Figure 5A).(12) We confirmed that these cells were capable of presenting *M. smegmatis*-derived antigen to MAIT cells upon induction of MR1A expression with doxycycline. Using this cell line, we could control the expression of MR1A expression by administering doxycycline either before or after transfecting the cells with pCI:MR1B_RFP (Figure 5B). To ensure that MR1AGFP expression was not artificially increased due to prolonged exposure to doxycycline, we showed that MR1AGFP expression remained steady at ∼86% GFP expression pre and post transfection with empty vector (Figure 6B, right). Interestingly, transfection of pCI:MR1BRFP into Beas2B:KO_Tet_MR1AGFP cells prior to the induction of MR1AGFP expression resulted in a 20% increase in MR1AGFP-negative cells, as compared to cells with doxycycline-induced MR1A expression prior to the transfection of pCI:MR1BRFP (Figure 5C). This indicated that the decreased MR1A protein abundance was the result of MR1B expression prior to induction of MR1A. In contrast, the expression of pCI: MR1CRFP either pre or post induction of MR1A had no effect on MR1A protein production (Figure 5C).

**Figure 5:**
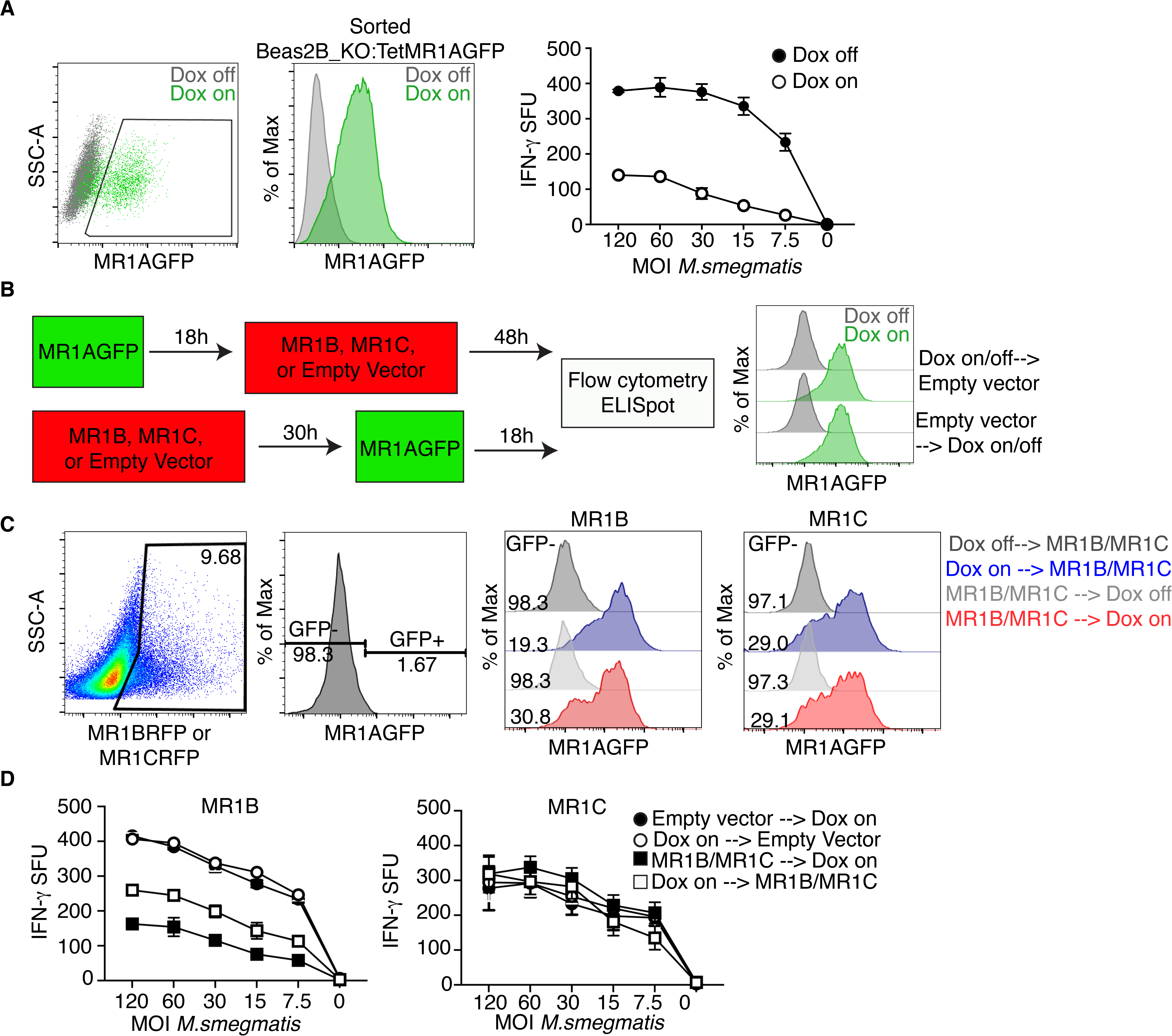
Prior MR1B expression decreases the abundance of MR1A protein. (A) Beas2B-MR1KO cells were transduced with a lentivirus encoding a doxycycline-inducible MR1AGFP and sorted based on MR1AGFP expression following overnight administration of doxycycline. Beas2B_MR1KO_TetMR1AGFP cells were treated with doxycycline overnight and infected with *M.smegmatis* at a titrating MOI for 1h. Infected cells were used as antigen presenting cells in an ELISpot assay as described earlier to stimulate MAIT production of IFN-*γ*. (B) Schematic of experiments to study the timing of MR1A and MR1B expression. Briefly, doxycycline was added to Beas2B_MR1KO_TetMR1AGFP cells prior to or following the transfection with MR1BRFP, MR1CRFP, or a pCI empty vector. MR1AGFP expression pre and post transfection with a pCI empty vector was measured by flow cytometry (right) (C) Transfected cells were assessed by flow cytometry, gated on MR1B or MR1C expression using RFP (left), and then assessed for GFP expression, which was based on a dox-off control (second). Overlay histograms of MR1AGFP expression following MR1B (third) or MR1C (fourth) transfection as described in Figure 6B with gMFI of MR1AGFP-negative condition reported. Doxycycline off transfection conditions are included as a control. Data are representative of at least 3 independent experiments with similar results. (D) ELISpot of cells transfected according to (B) and used as antigen presenting cells following infection with *M.smegmatis* at the indicated MOI. Cells were incubated with a MAIT clone and IFN-*γ* production was measured. Data are pooled from 3 independent experiments and error bars represent mean and standard error of replicates.

**Figure 6:**
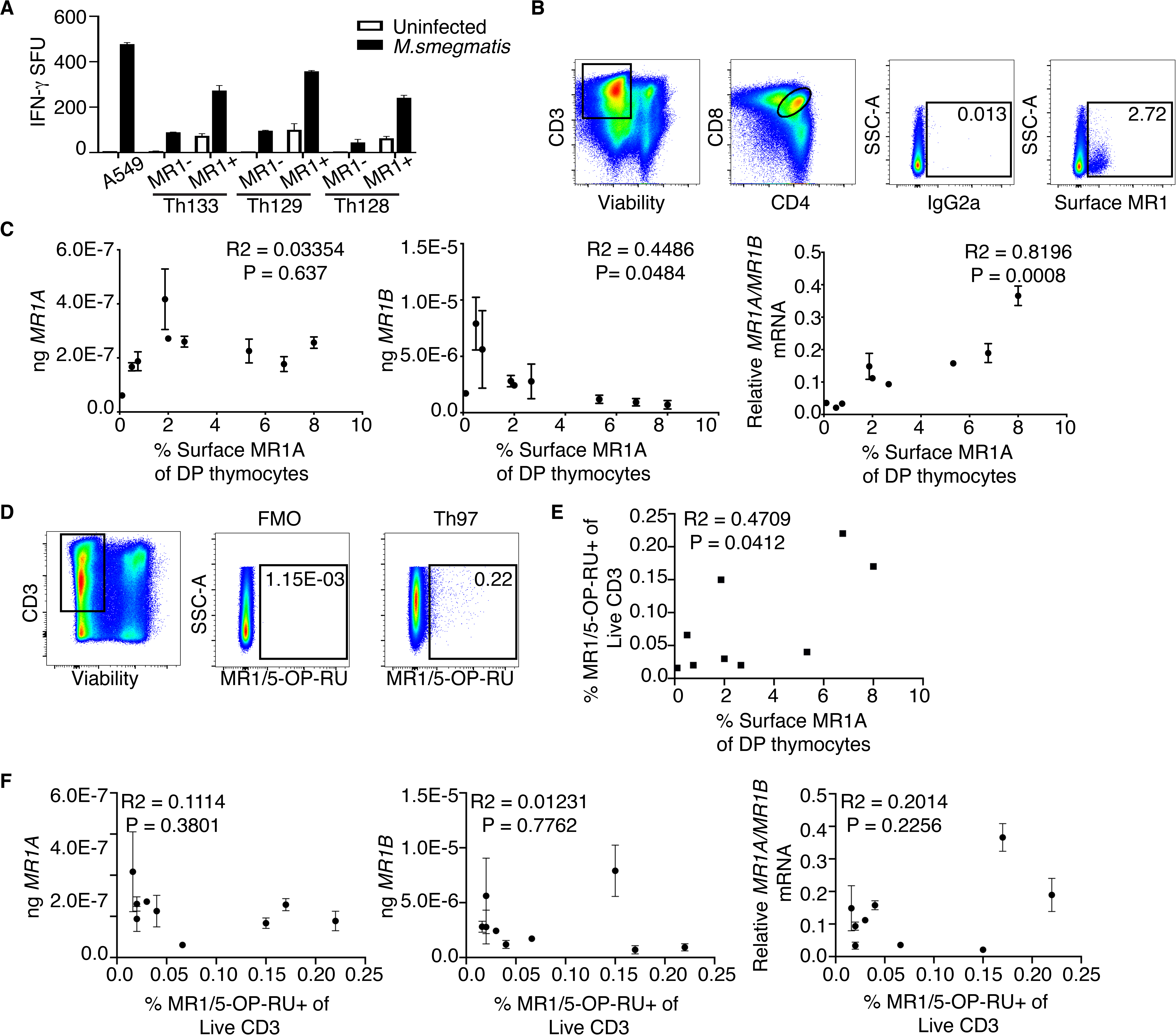
Double positive thymocyte expression of surface MR1A is associated with transcript expression and MAIT frequency in the thymus. MR1+ thymocytes were isolated using magnetic sorting based on MR1 expression (A) and used as antigen presenting cells following infection with *M.smegmatis* for 1h to stimulate MAIT production of IFN-*γ*, Error bars represent mean and standard error of duplicate wells. T cell production of IFN-*γ* is represented as spot forming units (SFU). (B) Gating strategy for sorting MR1+ CD4+ CD8+ (DP) thymocytes from human thymus. Cells were gated on live CD3+ CD4+ CD8+ MR1+ thymocytes based on an isotype control for MR1. (C) Absolute (left, middle), and relative (right) amounts of MR1A and MR1B transcript in DP MR1+ thymocytes versus surface expression of MR1 in 9 thymus donors. Pearson’s correlation was calculated and R^2^ and statistical significance is reported. (D) Gating strategy to identify MAIT cells in human thymus. Gates were set based on a negative control (FMO). Briefly, cells were stained with markers for viability, CD3, and the MR1/5-OPRU tetramer or the MR1/6-FP control tetramer (not shown). (E). Frequency of MR1/5-OPRU tetramer+ MAIT cells plotted versus surface MR1A expression in DP thymocytes from 9 donors. Pearson’s correlation was calculated and R^2^ and statistical significance is reported. (F) Same as (C), Expression of MR1A and MR1B transcript in DP thymocytes is plotted versus frequency of MR1/5-OPRU tetramer staining MAIT cells. Pearson’s correlation was calculated and R^2^ and statistical significance is reported.

To assess the impact of MR1BRFP expression prior to induction of MR1AGFP expression on the recognition of microbial ligands, we repeated the experiment as described in Figure 5B, and infected the cells with *M. smegmatis* (Figure 5D). We observed that, compared to the no-doxycycline control, Beas2B_MR1KO:Tet_MR1AGFP transfected with an empty vector were capable of stimulating MAIT cell activation upon the administration of doxycycline (Figure 5D). Upon transfection of MR1B in MR1A expressing cells, we observed that when MR1A and MR1B were both expressed, there was a 30% decrease in MR1A-mediated activation of MAIT cells, as seen in prior experiments with constitutively expressed MR1A. However, we observed an approximately 50% inhibition of MAIT activation when MR1B was expressed prior to induction of MR1A protein, corresponding to lower MR1A protein abundance under these conditions (Figure 5D, left). Expression of pCI:MR1CRFP pre or post induction of MR1A expression did not impact MR1A-mediated antigen presentation, indicating this was not a general consequence of transfection (Figure 5D, right). Thus, prior expression of MR1B may inhibit the abundance of MR1A protein, either through competing for chaperones necessary for MR1A folding, or by competing for factors required for protein synthesis of MR1.

### Expression of MR1 splice variants in CD4/CD8 MR1-expressing thymocytes

Cell surface expression of MR1 has been difficult to observe in the absence of either exogenous or bacterially-derived antigen(s).(31) We have previously shown that CD4+ CD8+ double positive (DP) thymocytes express high levels of MR1 on the cell surface.(32) The percentage of MR1-expressing thymocytes varies from 0.5% to 15% of DP thymocytes.(32) The role of MR1 expressing DP thymocytes in the selection of MAIT thymocytes is not well understood.(33) To determine whether MR1+ DP thymocytes are capable of presenting mycobacterial antigen to activate MAIT cells, we enriched for MR1-expressing DP thymocytes from human thymus using positive selection using magnetic beads following surface staining for MR1A. To determine if these cells could stimulate MAITs, the MR1+ enriched fraction as well as the MR1 depleted fraction were pulsed with *M.smegmatis* and tested for their ability to activate MAITs. As shown in Figure 6A, the MR1-expressing DP thymocytes could activate MAITs.

To assess whether relative expression of the MR1 isoforms is associated with prevalence of MR1A expressing thymocytes, we sorted MR1 expressing DP thymocytes from nine donors (Figure 6B). We then measured expression of *MR1A* and *MR1B* mRNA by qRT-PCR, which we then compared to surface MR1A expression in these donors (Figure 6C). We observe that there was not a strong association between *MR1A* mRNA and surface expression of MR1 (R^2^=0.03354, p=0.637), but there was a moderately significant association between *MR1B* transcript and the frequency of MR1+ DP thymocytes (R^2^=0.4486, p=0.0484). Intriguingly, we observed a strong positive association (R^2^=0.8196, p=0.0008) between relative *MR1A/MR1B* transcript and the frequency of MR1+ DP thymocytes.

To explore the relationship between DP MR1 expressing thymocytes and MAIT abundance in the thymus, we used the tetramer of MR1 bound to 5-OP-RU (MR1/5-OP-RU) to identify MAITs).(34) Among CD3 thymocytes, we observed frequencies of MAIT cells, with variation across donors (0.01-0.45% of live CD3+ cells). Intriguingly, the frequency of MR1 expressing thymocytes was associated with the frequency of MR1/5-OPRU+ thymocytes (R^2^= 0.5118, p=0.0302). Moreover, though the absolute amounts of *MR1A* and *MR1B* mRNA did not appear to be significantly correlated with frequency of MR1/5-OP-RU+ thymocytes, the ratio of *MR1A/MR1B* transcript in MR1-expression thymocytes was associated with the frequency of MAIT thymocytes (R=0.5501, p=0.0222) These results suggest a potential functional role for alternative splicing of MR1 in contributing to selection and development of MAIT cells in the thymus.

## Discussion

In this report, we show that MR1B, an alternatively spliced form of MR1, inhibits mycobacterial antigen presentation to MAIT cells by the full-length variant, MR1A. Both MR1A and MR1B are ubiquitously expressed, but with considerable variation between both discrete cell and tissue types and individual donors. These results suggest that MR1B could serve to modulate the recognition of MR1 ligands, and hence could play a role in the regulation of MAIT cell development and activation.

These results raise a number of questions about the mechanism(s) by which MR1B mediates this inhibitory function. As we found that MR1B did not traffic to the cell surface following exposure to exogenous ligand, we believe that MR1B is not directly blocking TCR engagement with MR1A. Additionally, as MR1A and MR1B reside in the same intracellular compartments, we postulate that the inhibitory effect of MR1B could occur in close proximity to the location of MR1A ligand loading. For example, MR1B may be competing for ligand, which either directly impedes the ability of MR1A to bind and present antigen to MAIT cells, or, blocks MR1A folding and stability. Alternatively, MR1B could compete for binding to a molecular chaperone which is necessary for the loading and or stabilization of the MR1A molecule following exposure to bacterial ligands, or for subsequent translocation of the loaded MR1A-ligand complex to the cell surface.

While coexpression of MR1A and MR1B resulted in decreased MAIT activation, we did not observe a decrease in MR1A protein abundance or surface expression. However, expression of MR1B prior to MR1A resulted in a decrease in total MR1A protein, further potentiating inhibition of antigen presentation. These data would support the concept that MR1B is interfering with access to ligand and/or chaperones that facilitate MR1A stability. Further studies involving analysis of MR1B protein structure would help shed light on whether MR1B can indeed bind ligand, as well as help elucidate the mechanism by which MR1B is acting to inhibit MR1A.

We found that relative expression of *MR1A and MR1B* varies considerably across tissues and donors. At present, the significance of this variation is not clear with regard to MAIT cell prevalence in tissues, their state of activation, and their ability to respond to MR1 antigens. Preliminarily, we note that tissues such as blood and spleen expressed relatively higher levels of *MR1A/MR1B*, while uterus, breast, and colon expressed lower *MR1A/MR1B* transcript. As these data were derived from mRNA from the whole tissue rather than specific cell types, the full significance of these data remains unclear. However, the wide variation would lead us to speculate that the relative abundance of MR1B could be associated with diminished activation of tissue resident MAITs. This could have significance with regard for the ability to recognize microbial infection or be associated with a propensity for autoimmunity.

Alternative splicing is often regulated by genetic modifications such as changes in DNA methylation, or, single nucleotide polymorphisms (SNPs).(35, 36) Recently, an intronic SNP in *MR1* was shown to be associated with susceptibility to tuberculosis.(37) These results raise the possibility that regulation of antigen presentation can be controlled at the level of gene expression. Alternately, we postulate that exposure to ligand or microbial infection may be playing a role in changing the expression of antigen presenting molecules, which may be key in understanding the regulation of antigen presentation by MR1.

Alternative splicing is estimated to occur at high frequency and represents an important means of controlling gene expression.(38, 39) Dysregulation in alternative splicing events has been associated with disease susceptibility, including those associated with autoimmunity and inflammation.(40) However, the role of alternative splicing in the regulation of MHC and subsequent acquisition of adaptive immunity is incompletely explored. An *α*3 domain deletion mutant of HLA-A11 was suggested to promote HIV evasion of NK cell mediated cytotoxicity.(41) Alternative splicing of the non-classical molecule HLA-G has been reported and is hypothesized to be associated with protection of the developing fetus.(42, 43) Additionally alternative splicing of MICA resulted in splice variants that were either agonists or antagonists for NKG2D.(44) These reports suggest that a putative role for alternative splicing is to control immune cell activation and therefore target immune responses to microbial infection. Analysis of disease cohorts, both in infection and autoimmunity, for differences in MR1 isoform expression would be helpful in understanding the physiological importance of alternative splicing of MR1.

Our results suggest a potential link between relative *MR1A/MR1B* expression and frequency of MR1A-expressing CD4+ CD8+ thymocytes. CD1d expressing DP thymocytes have been shown to contribute to the selection of iNKT cells, and MR1+ DP thymocytes could play a similar role in the selection of MAIT cells in the thymus.(45, 46) Staining of thymocytes with the MR1/5-OP-RU tetramer showed that the frequency of MAIT thymocytes was associated with the percentage of MR1A-expressing DP thymocytes. Notably, we did observe that while the frequency of MR1A expressing DP thymocytes varied, the overall protein levels of MR1A was relatively constant between donors. Therefore, it is clear that other factors are likely also are impacting the expression and/or stability of MR1A protein. We did observe that relative *MR1A/MR1B* expression on DP thymocytes was associated not only with frequency of MR1 expressing thymocytes, but also with the frequency of intrathymic MAIT cells. How DP MR1 expressing thymocytes function in MAIT development and the role of alternative splicing in this development will require detailed in vitro modeling. Furthermore, as we did not have matched PBMC from our thymus donors, we were not able to establish the relationship of intrathrymic MR1 expressing cells and the prevalence of MAITs in the blood.

Taken together, our results suggest that alternative splicing of MR1 represents a means of regulating and inhibiting MAIT activation and may be contributing to regulation of T cell responses to microbial antigen.

## Acknowledgements

We acknowledge the Pamela Canaday and the OHSU Flow Cytometry Shared Resource, Aurelie Snyder and the Advanced Light Microscopy Core Facility, as well as the Vollum DNA Sequencing Core for expert assistance. Lung and small intestine tissue were obtained from the Pacific Northwest Transplant Bank (PNTB). The gRNA to generate the MR1_KO cell line was a generous gift from Andrew Sewell (Cardiff University). Additionally, we thank Wilmon Grant for designing lentiviral constructs. We are grateful to Katherine Michaelis for critical reading of the manuscript. Finally, we thank Christopher Wilks for help using Snaptron.

